# Transcriptome responses of intestinal epithelial cells induced by membrane vesicles of *Listeria monocytogenes* unveil novel insights into the host-pathogen cross talk

**DOI:** 10.1101/679951

**Authors:** Raman Karthikeyan, Pratapa Gayathri, Paramasamy Gunasekaran, Medicharla V. Jagannadham, Jeyaprakash Rajendhran

**Author notes:** Address correspondence to Jeyaprakash Rajendhran, and Medicharla V. Jagannadham, (.).

## Abstract

Membrane vesicles (MVs) serve as a vital source of virulence factors in many pathogenic organisms. The release of MVs by *Listeria monocytogenes* is only recently recognized, but its role in the pathogenesis is poorly understood. Here, we investigated the role of MVs of *L. monocytogenes* in virulence and host interactions. Proteomic analyses of whole cells and MVs of *L. monocytogenes* were performed using LC/MS/MS. A total of 1376 and 456 proteins were identified in the *L. monocytogenes* cells and MVs, respectively. Also, we have found that MVs contains active pore-forming listeriolysin (LLO), internalin B (inlB), phosphatidylinositol-specific phospholipase C (PI-PLC-A). We have previously reported that MVs of *L. monocytogenes* can infect and induce cytotoxicity in Caco-2 cells. In this study, we report the transcriptome response of Caco-2 cells upon infection with MVs as well as *L. monocytogenes*. In particular, we observed the up-regulation of autophagy-related genes in the early phase of infection with MVs. Transcription of inflammatory cytokines (CCL2, CXCL6, CXCL8, CXCL15, CXCL5, CXCL10) peaked at four h of infection. A large number of differentially expressed genes was associated with actin cytoskeleton rearrangement, autophagy, cell cycle arrest, and induction of oxidative stress. At a later time point, transcriptional programs generated upon infection with MVs point toward to evade innate immune signals, by modulating the expression of anti-inflammatory genes. KEGG pathway enrichment analysis revealed that MVs induce several signaling pathways such as PI3k-Akt signaling pathway, mitogen-activated protein kinase (MAPK) pathway, NOD-like receptor signaling pathway, cAMP signaling pathway, TNF, and NF-kB signaling pathway. Moreover, MVs induced the expression of cell cycle regulatory genes, which may result in the ability to prolong host cell survival, thus protecting the replicative niche for *L. monocytogenes*. Notably, we identified several non-coding RNAs (ncRNAs) are regulated during infection, suggesting that an early manipulation of the host gene expression may be essential for *L. monocytogenes* persistence and replication in host cells.

## Introduction

Several Gram-negative and Gram-positive bacteria release membrane vesicles (MVs). MVs are a spherical structure of 50-300 nm in diameter composed of the single membrane that is derived from the outer membrane (OM) and contains OM proteins, lipopolysaccharide (LPS), and other lipids, while the vesicle lumen mainly contains periplasmic proteins (1). Many pathogenic Gram-negative bacteria species extend their virulence potential by releasing spherical buds, derived from the outer membrane, so-called outer membrane vesicles (2). The release of MVs can benefit the microbe, mediating microbial interactions with the human host and within bacterial communities (3,2). Also, MV production protects bacteria under stressful conditions by eliminating the accumulated damaged DNAs and proteins (4). Functions ascribed to MVs include the promotion of virulence, biofilm formation, signal transduction, cytotoxicity, and host pathology (5,6). Also, MVs contains various cargo molecules which can modulate host pathology (7–9). Though MVs of Gram-negative bacteria have been extensively studied, MVs of Gram-positive bacteria were overlooked for decades and yet to be explored. In recent years, several Gram-positive bacteria such as *Staphylococcus aureus*, *Streptococcus* spp., *Clostridium perfringens*, and *Listeria monocytogenes* have been reported to produce membrane vesicles (MVs) (10–14).

*L. monocytogenes*, the etiologic agent of listeriosis, remains a serious public health concern with its frequent occurrence in food coupled with a high mortality rate (15,16). The success of *L. monocytogenes* results from the ability to promote its internalization by host cells, which enables the bacterium to overcome the host defense mechanisms and to survive inside the host cells. *L. monocytogenes* use an array of virulence effectors that act in one or more steps during the cellular infection (17). The MVs can serve as a safe delivery vehicle for contact-independent targeted delivery of toxins to the host cells. Upon delivery of virulence factors, MVs perform an essential function in bacterial pathogenesis without the direct interaction between pathogen and host cell (18,8). In our previous study, we reported the proteome analysis of *L. monocytogenes* MTCC 1143 (Serotype 4b) (19). A total of 16 unique proteins were identified only in MVs, including PI-PLC, autolysin, uncharacterized protein *yabE,* competence protein ComEC/Rec2, flagellar proteins, and other uncharacterized proteins. These proteins have essential roles in virulence, adaptation, metabolism, and regulation.

Several studies have investigated the host cell responses to the *L. monocytogenes* infection (20), whereas there is a lack of information on the host cell responses to its MVs. Here, we report the host cell responses to the MVs of *L. monocytogenes*. RNA-Seq technology provides a powerful approach for studying global response during infection (21,22). RNA-Seq allows the simultaneous sequencing of millions of RNA transcripts, thus enabling identification of global gene expression under different conditions (23,24). Here, we have used RNA-Seq to map the changes in host transcriptome during MV infection globally. Differential gene expression analysis revealed the modulation of gene expression over time; pathway and gene ontology analyses provided novel insights into the processes during infection.

We focused on host cell responses during the early phase of interaction, as this period is essential for the establishment of infection. We identified several pathways that are up-regulated during infection and are central for *Listeria* persistence. We identified MV-mediated signaling pathways and significant dysregulation of the microtubule and cytoskeletal network in the host cell by MVs. Also, a set of novel small nucleolar RNAs (snoRNAs), micro-RNAs (miRNAs), and small Cajal body-specific RNAs (scaRNAs) were identified that could represent novel biomarkers of *L. monocytogenes*-induced infections. Also, pathways down-regulated during the MV infection were mapped, which include the cell adhesion, cytoskeletal regulators and apoptosis, and extracellular matrix synthesis. Our data represent the first transcriptome analysis of dynamic interaction of MVs with intestinal epithelium and provide new insights into the MV-mediated pathogenesis of *L. monocytogenes*.

## Materials and Methods

### Bacterial strains and culture growth

*L. monocytogenes* 10403S strain was routinely grown on brain heart infusion agar (BHI) (Himedia; Mumbai India) at 37°C.

### Proteome analysis

Extraction of whole cell proteome, isolation, and purification of MVs, dynamic light scattering (DLS) analysis, transmission electron microscopy, SDS-PAGE and in-gel digestion, LC/MS/MS and bioinformatic analysis were done as described in our previous paper (19).

### Caco-2 cell infection by *L. monocytogenes* and MVs and RNA extraction

Growth and maintenance of the human colon adenocarcinoma cell line, Caco-2, were done as described earlier (19). For bacterial infection, Caco-2 cells in antibiotic-free cell culture media were seeded in a 6-well plate (Hi-media) at a concentration of 1 × 10^5^ cells. Cells were then infected in triplicate with *L. monocytogenes* 10403S at a multiplicity of infection (MOI) of 10 and incubated at 37°C, 5% CO_2_. After 1 h infection period, infected monolayers were washed once with 1 ml sterile PBS, and gentamicin was added (50 µg/ml) at this point. After, the cells were incubated for further time points (4 h and 8 h) at 37°C, 5% CO_2_. Similarly, for MV infection, Caco-2 cells were treated with appropriate concentrations of MVs and incubated for 4 h and 8 h at 37 °C, 5% CO_2_. Uninfected cells were used as control. After infection, at each times points, cells were washed and scrapped off gently. Total RNA was then extracted using the Qiagen RNeasy mini kit according to the manufacturer’s instructions. The concentration and quality of extracted RNA were assessed using an Agilent 2100 bioanalyzer and stored at − 80 °C.

### Library preparation, RNA sequencing, and data analysis

Illumina NextSeq RNA Sample Prep Kit (version 3) was used according to the manufacturer’s instructions for RNA-Seq sample preparation. Poly (A)-enriched cDNA libraries were generated using the Illumina NextSeq sample preparation kit (San Diego, CA) and checked for quality and quantity using the Bioanalyzer. Paired-end reads were obtained using the Illumina NexSeq 500 platform. Trimmomatic was used to remove any remaining Illumina adapter sequences from reads and to trim bases off the start or the end of a read when the quality score fell below a threshold (25). Sequence quality metrics were assessed using FastQC. Reads were aligned independently to the *Homo sapiens* genome (GRCh38/hg38) obtained from the UCSC genome browser (http://genome.ucsc.edu) using HISAT (v 2.0.10) (26). After read mapping to hg38, SAMtools (http://www.htslib.org/) was used to filter bam files for uniquely mapped reads. Resulting BAM files were used for all further downstream analyses.

Reads aligned to annotated genes were quantified with the feature-count program Subread (27) using gene annotations from the UCSC genome browser. Uniquely mapped reads were subjected to DEseq2-Bioconductor R package (28) for identification and quantification of genes that were significantly differentially expressed between the conditions, following standard normalization procedures. The FPKM (fragment per kilobase max. transcript length per million mapped reads) values were computed for each library from the raw read counts. Normalized counts were transformed to rlog values to create a heat map. The list of DESeq2 determined differentially expressed genes (DEGs) was filtered with a conservative absolute log2 fold change cutoff and corrected *p*-value (p<0.05). Heatmap was generated using Heatmapper (29). Differentially expressed genes of data sets were subjected to the enhanced volcano to create volcano plot (https://github.com/kevinblighe/EnhancedVolcano). Boxplot was generated using BoxPlotR (30). Lists of differentially expressed genes were further annotated with ConsensusPathDB (31), BionetDB (32) and pathway information from the KEGG database.

### Gene ontology (GO) enrichment analysis

GO categories enriched in the DE gene lists were identified using the ConsensusPathDB (31) and DAVID (33). For each comparison, up-regulated and down-regulated gene sets were subjected to ConsensusPathDB and DAVID. A P value cut-off of 0.01 was used.

### KEGG pathways and network analyses

KEGG pathway analysis using ConsensusPathDB-*Homo sapiens* was done to identify signaling and metabolic pathways that were over-represented in the DE gene lists. For each KEGG pathway, a P value was calculated using a hypergeometric test, and a cutoff of 0.01 was applied to identify enriched KEGG pathways. Genes that were more than 2-fold in infected cells relative to uninfected controls at each time point were used as an input with up- and down-regulated genes considered separately.

### Non-coding RNA analysis

Noncoding RNA and microRNA were predicted from lncRNome (http://genome.igib.res.in/lncRNome/), LNCipedia (www.lncipedia.org), and miRBase (www.mirbase.org) databases.

## Results

### Purification of MVs from *L. monocytogenes* strain 10403S

*L. monocytogenes* was grown in BHI broth and MVs were isolated from culture supernatant by ultracentrifugation. The electron micrographs revealed that the MVs of *L. monocytogenes* are spherical with a uniform size distribution. The mean hydrodynamic radius of vesicles was 192.3 nm as determined by DLS. All further experiments were performed using these purified MVs.

### MV proteome of *L. monocytogenes* 10403S

*L. monocytogenes* cells and density gradient fractions (60%) were resolved by SDS-PAGE to visualize the MV associated proteins (Fig. 1). The proteome of the *L. monocytogenes* cells and purified MVs were analyzed by LC-MS/MS. A total of six LC-MS/MS runs, three each for cellular proteome and MV proteome were carried out. Based on SEQUEST and PEAKS analysis, 8,007 and 1,437 high confident peptides were identified in *L. monocytogenes* cells and MVs, respectively. The complete list of the identified peptides and proteins are provided in Table S1 and S2, respectively. From these peptides, 1376 and 456 proteins were identified in *L. monocytogenes* cells and MVs, respectively. We compared the MV and cellular proteomes of *L. monocytogenes*. As shown in the Venn diagram, 401 proteins were found in both whole cell and MV proteome (Fig. 1B). Among the 456 proteins in MVs, 235 proteins (52%) were predicted as cytoplasmic, 63 proteins (14%) as cytoplasmic membrane, 42 proteins (9%) as extracellular, 11 proteins (2%) as cell well-associated. Localization of 105 proteins (23%) could not be predicted using Psortb (Fig.1C and D).

**Fig.1.**
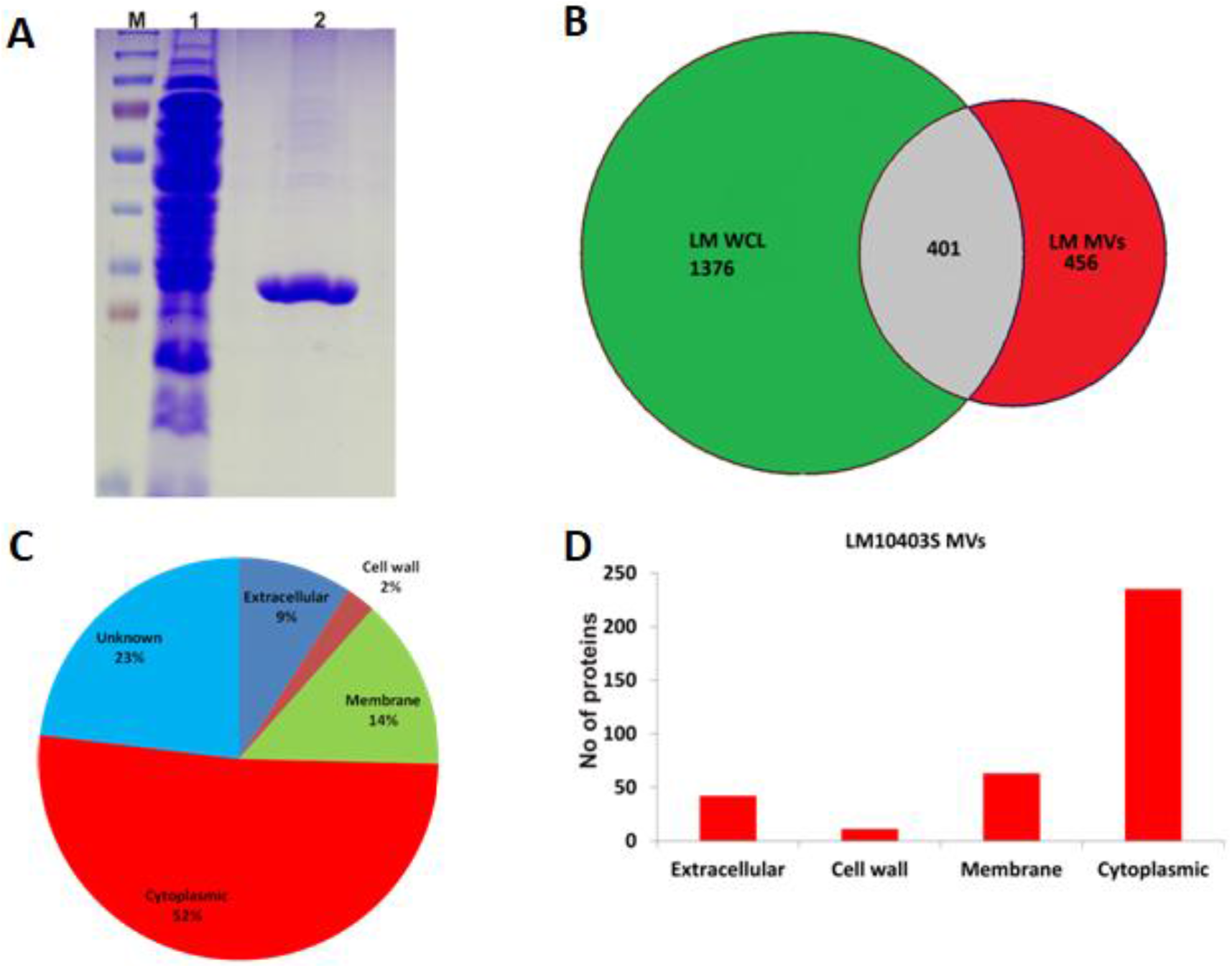
Proteomic characterization of MVs of *L. monocytogenes*. (A) *L. monocytogenes* cells and purified MV proteins were resolved by SDS-PAGE. The gels shown are the actual SDS-PAGE lanes that were analyzed. (B) The Venn diagram describes individual differences in the total number of proteins identified exclusively in MVs (but, not in whole-cell proteome). (C & D) Sub-cellular location of MV-associated proteins. The proteins identified through LC-MS/MS profiling were processed using PSORTb and CELLO. The MV profiles that were predicted to be in the cytoplasm, membrane, extracellular, and cell wall were counted, and their percentage is depicted.

Among the identified MV proteins, few posses LPXTG motif, which is known to mediate the covalent attachment to the cell wall and a few carry LysM motif that allows non-covalent attachment to the peptidoglycan. A total of 40 unique proteins were found only in MVs such as, oligopeptide transport ATP-binding protein oppF, ABC-2 type transport system permease, chromosomal replication initiator protein DnaA, DNA mismatch repair protein MutL, DNA segregation ATPase FtsK/SpoIIIE, flagellar protein export ATPase FliI, transcriptional pleiotropic regulator, secreted protein and few other uncharacterized proteins (Table 1).

**Table.1.**
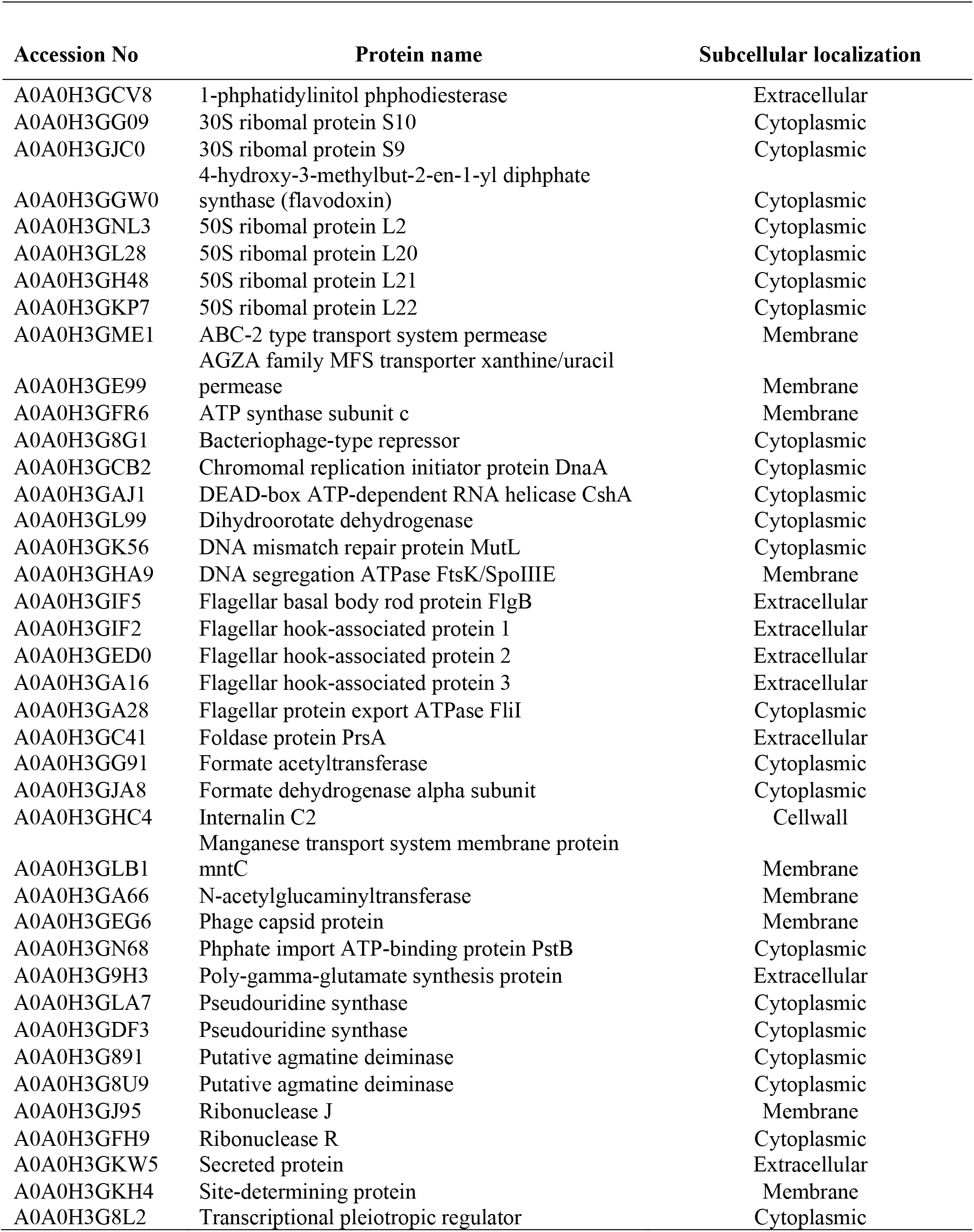
List of proteins identified only in MVs.

### Virulence factors associated with MVs

Major virulence factors such as LLO, InlB, PLC-A, flagellin, HlyD family secretion protein, LemA protein, methyl-accepting chemotaxis protein, N-acetylmuramoyl-L-alanine amidase, and pheromone lipoprotein were identified in MVs. Also, we have identified major secretory proteins prsA, invasion associated secreted endopeptidase, and multifunctional fusion protein (secD), murA, and SecA in MVs (Table 2). Also, sortase A was identified in MVs with the LPXTG motif, known to mediate the covalent attachment of the protein to the cell wall. The role of sortase A in *L. monocytogenes* infection is critical as several effectors were found to be associated with the bacterial surface through the activity of this enzyme.

**Table.2.**
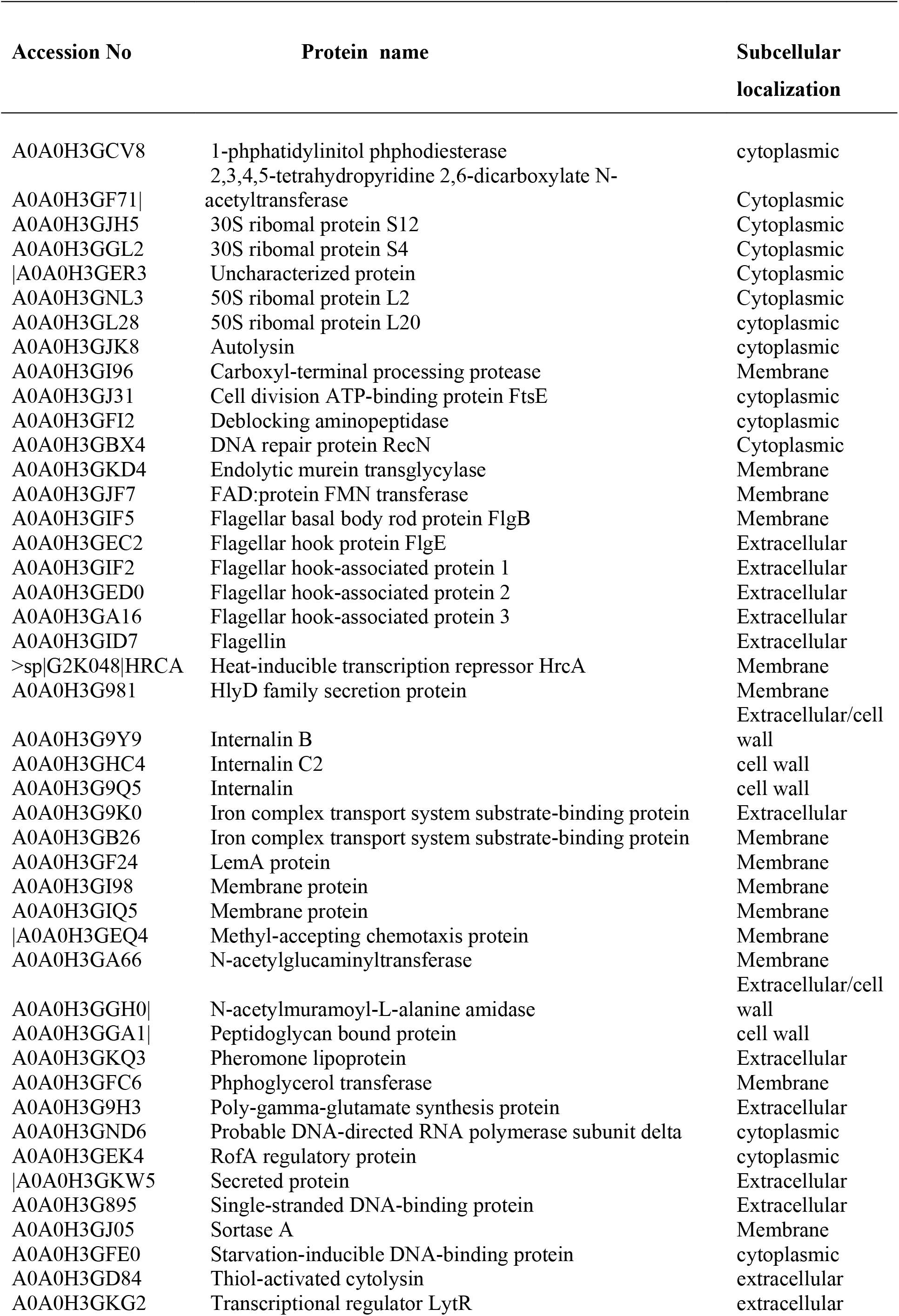
List of virulence proteins identified in MVs.

### *L. monocytogenes* MVs constitutes of lipoproteins

Lipoproteins have been described as virulence factors because they play critical roles in virulence, membrane stabilization, nutrient uptake, antibiotic resistance, bacterial adhesion to host cells, and secretion and many of them still have unknown function. We have identified 30 lipoproteins in *L. monocytogenes* MVs including pheromone lipoprotein, manganese-binding lipoprotein mntA, membrane protein insertase YidC, and several other uncharacterized lipoproteins.

### Chaperones associated with *L. monocytogenes* MVs

A specific group of ATP-dependent chaperones, called foldases, assist the correct post-translocational folding of the secreted proteins. PrsA, a ubiquitous Gram-positive lipoprotein with peptidyl-prolyl cistrans isomerase activity, was identified in MVs. Also, we have identified chaperon proteases, heat-inducible transcription repressor HrcA, GroS, GroL, HslO.

### Transporters

We have identified several transporter proteins in MVs, including carnitine transport ATP-binding proteins such as OpuCA, OpuCC. The osmolyte transporters involved in the uptake of glycine, betaine, BetL, and Gbu, were identified in MVs. Also, divalent metal cation transporter MntH, oligopeptide transport ATP-binding protein oppF and ABC transporters were detected in MVs.

### MV infection of intestinal epithelial cells extensively altered the host transcriptome

Caco-2 cells were treated with MVs and incubated for 4 h and 8 h post-challenge in three biologically independent experiments per time point. At 4 h post-infection, a total of 1,189 transcripts were significantly altered upon infection of Caco-2 cells with *L. monocytogenes* (>1.5- fold regulation, p-value<0.05, p-adj<0.05); of these, 669 were up-regulated and 520 were down-regulated. At 8 h post-infection, 989 genes were significantly altered. Of these, 360 were up-regulated while 629 were down-regulated (>1.5- fold regulation, pvalue<0.05, padj<0.05). MA plot was generated for all the data sets, and shown in Fig. 2. Differentially expressed genes (DE) in infected cells compared to the untreated cells were identified by calculating reads per Fragment per kilobase per million (FPKM). The top 50 up- regulated and down-regulated genes are shown in Table. 3. Gene set enrichment analysis using ConsensusDB and DAVID identified significantly enriched functional groups that were altered by MV infection.

**Table 3.** List of top 50 up-regulated genes upon infection with MV 4h.

| **RefSeq mRNA**<br>**Accession** | **Gene Symbol** | **fold change** | **pvalue** |
| --- | --- | --- | --- |
| NM_001013257 | BCAM | 183.827064 | 2.80E-08 |
| XM_017001418 | XM_017001418 | 74.21906482 | 1.10E-05 |
| NR_027011 | YBX3P1 | 65.04272479 | 1.17E-10 |
| NM_000558 | HBA1 | 63.37685609 | 0.000323858 |
| NR_135053 | LOC101928424 | 62.51044099 | 0.00024127 |
| XM_017018991 | ETV6 | 62.26420224 | 0.000157274 |
| NM_002982 | CCL2 | 59.72644989 | 0.000215706 |
| XM_011528501 | TOMM34 | 57.9153817 | 0.007545992 |
| XR_001746888 | LOC107987113 | 57.28190218 | 0.00019115 |
| XR_924707 | LOC102724479 | 57.03544064 | 0.000613036 |
| NM_001172896 | CAV1 | 55.56831349 | 0.000233717 |
| NM_005447 | RASSF9 | 54.5221971 | 0.000298389 |
| NM_030762 | BHLHE41 | 52.702984 | 0.00051697 |
| NM_152337 | C16orf46 | 51.12359277 | 0.00041761 |
| NM_000584 | CXCL8 | 50.23243015 | 3.98E-21 |
| NR_046748 | ARHGAP31-AS1 | 47.85203229 | 0.000965889 |
| XR_001748366 | XR_001748366 | 46.0395765 | 0.001359089 |
| NM_001190484 | CYP3A5 | 43.57338401 | 1.34E-06 |
| NM_002994 | CXCL5 | 42.61733486 | 1.15E-05 |
| XM_017029555 | OPHN1 | 41.73769274 | 0.000300671 |
| NM_001293626 | MGAM2 | 40.10182622 | 3.00E-05 |
| NR_110840 | LOC101927136 | 38.56446007 | 0.000593169 |
| NM_001127380 | SAA2 | 33.88647895 | 6.95E-06 |
| XR_932500 | LOC105370910 | 33.34294695 | 0.001277114 |
| NM_018256 | WDR12 | 31.65464133 | 1.69E-14 |
| XM_017012837 | RAPGEF5 | 30.73065253 | 0.001416844 |
| XM_017011596 | LOC107986532 | 30.71038328 | 0.001284758 |
| NM_022549 | FEZ1 | 29.43280071 | 8.21E-06 |
| XM_005271963 | AFF4 | 29.11498113 | 0.002510524 |
| XM_011511452 | XM_011511452 | 28.86791012 | 0.000362025 |
| NM_002421 | MMP1 | 28.055432 | 1.30E-12 |
| NR_002941 | PDCL3P4 | 26.99787104 | 0.000421435 |
| XM_005270920 | XM_005270920 | 26.86905861 | 0.004509202 |
| NM_000612 | IGF2 | 25.39494254 | 7.82E-12 |
| NM_002427 | MMP13 | 24.73422212 | 0.003630716 |
| NM_024093 | C2orf49 | 23.79018336 | 3.30E-11 |
| XM_017004977 | CAMKMT | 23.69013751 | 0.0046411 |
| XM_017018460 | TRIM5 | 23.58165051 | 0.0061141 |
| NM_001565 | CXCL10 | 23.09092898 | 3.87E-11 |
| NM_139284 | LGI4 | 22.34027104 | 9.70E-07 |
| XR_926322 | ADTRP | 22.14151853 | 0.007547942 |
| XR_001749448 | LOC107984461 | 21.89489511 | 0.006535032 |
| XR_925333 | TBC1D19 | 20.04499122 | 0.000120534 |
| XR_001746017 | LOC107986959 | 19.7562818 | 0.001740063 |
| XR_944886 | LOC105369743 | 19.71404729 | 0.000333113 |
| NM_025047 | ARL14 | 19.06105327 | 7.22E-16 |
| NM_001164431 | ARHGAP40 | 17.81523553 | 0.000297346 |
| NM_001039141 | TRIOBP | 17.47158308 | 0.003333792 |
| NM_001318000 | CWC27 | 17.02771847 | 0.00469703 |

**List of top 50 down-regulated genes upon infection with MV 4h**
| **RefSeq mRNA**<br>**Accession** | **Gene Symbol** | **fold change** | **pvalue** |
| --- | --- | --- | --- |
| NR_024230 | SNAR-B2 | 0.030131 | 0.007254 |
| NR_132775 | SNORA93 | 0.028662 | 0.000968 |
| XM_017019183 | ANKRD52 | 0.028475 | 0.000919 |
| NR_003021 | SNORA79 | 0.028144 | 0.000959 |
| XM_005248038 | TMEM129 | 0.027943 | 0.000972 |
| NR_003944 | SNORD78 | 0.027906 | 1.30E-09 |
| NM_003740 | KCNK5 | 0.027387 | 0.001085 |
| NM_001012716 | TYMSOS | 0.026715 | 3.81E-05 |
| NM_024036 | LRFN4 | 0.026029 | 2.52E-08 |
| NR_003035 | SNORA16A | 0.024895 | 9.07E-21 |
| NM_006452 | PAICS | 0.024778 | 0.000447 |
| NR_004392 | RNY3 | 0.023652 | 2.58E-05 |
| NR_006880 | SNORD3A | 0.023581 | 6.73E-21 |
| NR_024231 | SNAR-B1 | 0.0215 | 0.004253 |
| XM_011538197 | ANKRD52 | 0.021156 | 0.00017 |
| NR_104085 | RNVU1-6 | 0.021059 | 2.79E-05 |
| NR_002736 | SNORD60 | 0.01985 | 5.77E-14 |
| NM_030782 | CLPTM1L | 0.019748 | 9.86E-17 |
| NR_003941 | SNORD75 | 0.019362 | 7.07E-10 |
| NR_002979 | SNORA49 | 0.019156 | 5.82E-07 |
| NR_130740 | FAM86EP | 0.019075 | 0.000131 |
| NM_213611 | SLC25A3 | 0.019051 | 0.000185 |
| NR_002952 | SNORA9 | 0.018407 | 2.46E-25 |
| NR_003271 | SNORD3B-1 | 0.018112 | 1.59E-27 |
| XR_926571 | LOC105374968 | 0.01807 | 0.000441 |
| NR_000008 | SNORD22 | 0.018046 | 1.92E-06 |
| NR_003019 | SNORA77 | 0.01711 | 0.000421 |
| NM_001098208 | HNRNPF | 0.017105 | 0.000359 |
| NR_003706 | SNORA38B | 0.017081 | 6.74E-07 |
| NR_003018 | SNORA71D | 0.016877 | 1.93E-22 |
| XM_017024762 | XM_017024762 | 0.01581 | 0.000217 |
| NR_003017 | SNORA71C | 0.015122 | 2.47E-05 |
| NR_002602 | SNORD37 | 0.014539 | 0.000114 |
| XM_011544161 | PGBD2 | 0.014519 | 0.000108 |
| XR_001754568 | XR_001754568 | 0.013777 | 8.12E-05 |
| NR_004378 | SNORD94 | 0.013696 | 7.80E-23 |
| NR_003020 | SNORA78 | 0.013122 | 6.02E-05 |
| NR_004403 | SNORD97 | 0.012653 | 5.03E-52 |
| NM_000112 | SLC26A2 | 0.012236 | 5.33E-06 |
| NR_002911 | SNORA71A | 0.011294 | 3.14E-27 |
| NR_002604 | SNORD10 | 0.010325 | 0.000464 |
| NR_003048 | SNORD23 | 0.010241 | 1.67E-05 |
| NR_002918 | SNORA48 | 0.009871 | 2.54E-06 |
| NR_109922 | LAMA5-AS1 | 0.008899 | 4.57E-06 |
| NR_002999 | SCARNA20 | 0.005176 | 5.94E-09 |
| NM_001032 | RPS29 | 0.005039 | 1.94E-37 |
| NR_024225 | SNAR-A11 | 0.003089 | 2.84E-10 |
| NR_004436 | SNAR-A2 | 0.002974 | 1.59E-10 |
| NR_031639 | MIR1304 | 0.002158 | 8.29E-12 |

**Fig.2.**
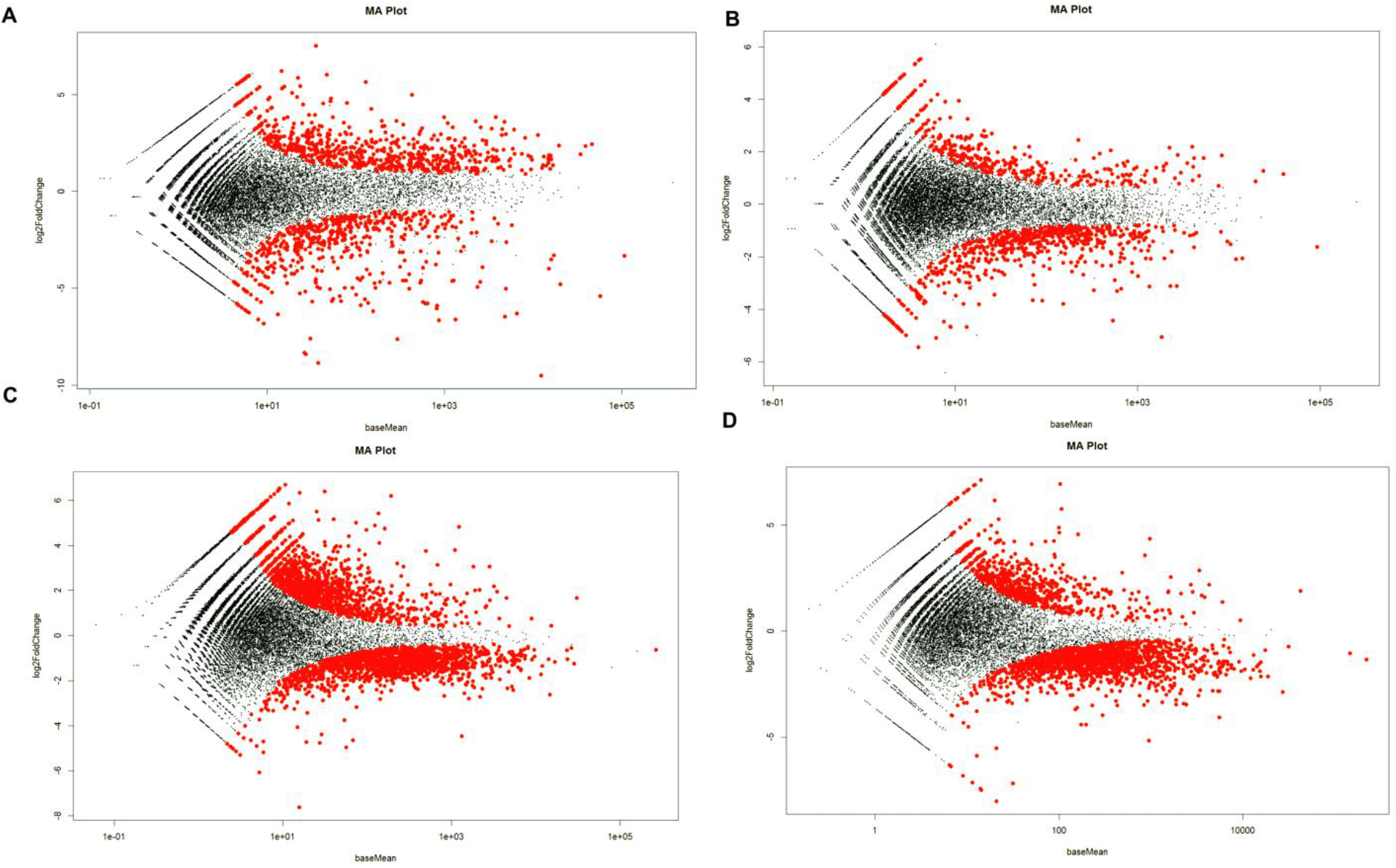
The host transcriptional response to the infection by *L. monocytogenes* and its MVs. (A) Bland-Altman (MA) plots showing the differential expression of genes in Caco-2 cells after 4 h of infection with MVs compared to the control. (B) at 8 h of infection with MV (C) at 4 h infection with *L. monocytogenes* (D) at 8 h infection with *L. monocytogenes*. The plots were generated by DESeq R package.

### Epithelial response to infection with MVs

Primary response genes, which include immediate-early and delayed genes, play pivotal roles in a wide range of biological processes including differentiation, proliferation, survival, stress, innate and adaptive immune responses, and glucose metabolism. At 4 h, many of the up-regulated genes belong to endocytosis, autophagy, actin cytoskeleton rearrangements, cell cycle, and pro-inflammatory cytokines. Genes were categorized according to biological function and canonical pathways. The top GO terms enriched in each category with the p-values are shown in Fig 3.

**Fig.3.**
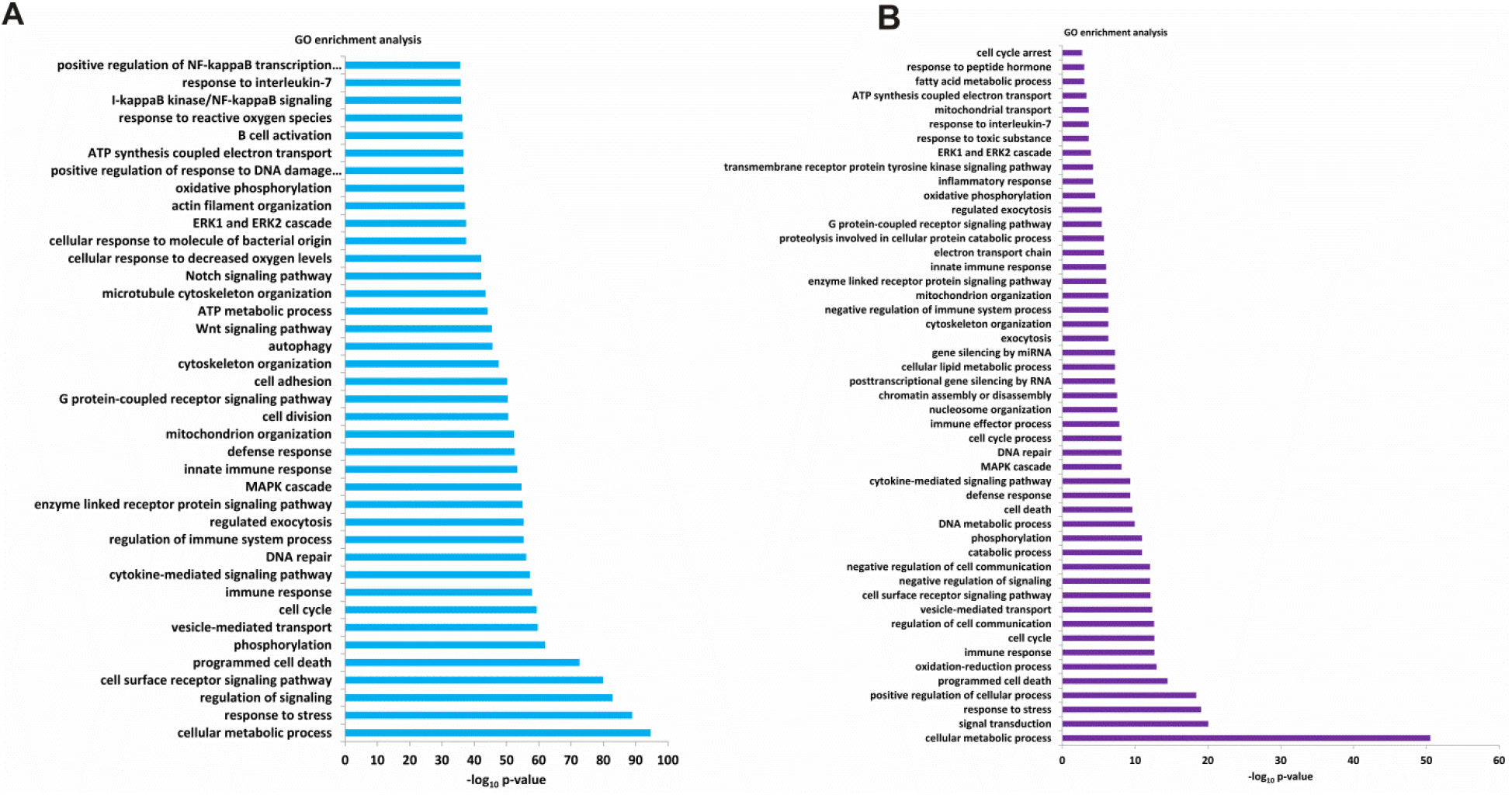
Gene ontology enriched terms for genes differentially regulated in Caco-2 cells upon infection with MVs. (A) Pathways significantly regulated in intestinal epithelial Caco-2 cells infected with MV 4h (B) Pathways and process significantly regulated in intestinal epithelial Caco-2 cells exposed to MVs at 8h infection. Terms were sorted according to q-value, and FDR corrected P-value.

The GO enrichment analysis in biological processes revealed the unique features of MV infection. The most enriched early up-regulated clusters include the endocytosis, defense response, and pro-inflammatory response. In contrast, the most enriched early down-regulated group is involved in the regulation of metabolic pathways. Gene expressions of cellular pathways were noticeably altered by infection. Majority of the affected pathways are vital for innate immune responses (*P* < 0.05, tested by Wilcox test, followed by the Benjamini and Hochberg multiple test correction), according to the Kyoto Encyclopedia of Genes and Genomes (KEGG, http://www.genome.jp/kegg/) annotation (Fig. 4). Invading microbial pathogens are sensed by the host PRR, such as Toll-like receptors (TLR), NOD-like receptors (NLR), and RIG-I-like receptors (RLR), through binding of PRRs to their ligands called pathogen-associated molecular patterns (PAMPs), which leads to the activation of host immune responses to microbial infections. DEGS in TLR and NLR signaling pathways, as well as cytokine-cytokine receptor interaction and chemokine signaling pathways, were significantly enriched (Table 4). Most of the pathways associated with innate immune responses were up-regulated, whereas those involved in the basal cellular metabolic pathways were generally down-regulated.

**Table.4.** KEGG pathways enriched in MV infected cells.

| KEGG term | Description | Genes | P-value |
| --- | --- | --- | --- |
| hsa04668 | TNF signalling pathway | CXCL10, CFLAR, CXCL5, CXCL2, CXCL3, PIK3CA, CCL2, JUN, PTGS2, CREB3L1, MMP14, ICAM1, NFKBIA | 0.00024414 |
| hsa04621 | NOD-like receptor signalling pathway | CXCL2, GBP1, GBP2, CCL2, JUN, BRCC3, NFKBIA, TICAM1, CXCL3, CXCL8, NFKBIB, GABARAPL1 | 0.000488 |
| hsa04657 | IL-17 signalling pathway | MMP13, CXCL10, CXCL5, CXCL2, CXCL3, CCL2, JUN, PTGS2, HSP90B1, MMP1, CXCL8, NFKBIA | 0.000488 |
| hsa04062 | Chemokine signaling pathway | CXCL10, CXCL5, CXCL2, CXCL3, PIK3CA, CCL2, CXCL16, NFKB, IGNG5, CXCL8, NFKBIB, SOS1, SOS2 | 0.001709 |
| hsa05164 | Influenza A | TMPRSS13, DDX58, CXCL10, HSPA1A, TMPRSS4, PIK3CA, CCL2, JUN, NFKBIA, NXT1, TLR7, TICAM1, ICAM1, CXCL8, NFKBIB, HSPA2 | 0.002686 |
| hsa04064 | NF-kappa B signaling pathway | CSNK2A3, CFLAR, CXCL2, DDX58, PTGS2, NFKBIA, TICAM1, ICAM1, CXCL8 | 0.003906 |
| hsa05160 | Hepatitis C | DDX58, EGFR, PIK3CA, NFKBIA, CLDN, CLDN3, TICAM1, CXCL8, EIF3E, SOS1, SOS2 | 0.004883 |
| hsa05200 | Pathways in cancer | TXNRD1, MYC, TPR, MET, JUN, HSP90B1, IGF2, CALM1, EGFR, PIK3CA, RALBP1, NFKBIA, SHH, LAMC1, E2F1, LAMC2, MMP1, CCDC6, GNG5, IL2RG, PTGS2, PMAIP1, CXCL8, RBX1, CDK4, SOS1, SOS2 | 0.006998 |
| hsa05134 | Legionellosis | HSPA1A, CXCL2, CXCL3, HSPA2, C3, NFKBIA, CXCL8, EEF1A1, EEF1A2 | 0.007813 |
| hsa04010 | MAPK signaling pathway | MYC, EGFR, HSPA1A, MET, HSPA2, JUN, CACNG8, CACNG4, EREG, AREG, DUSP5, DUSP6, IGF2, NF1, SOS1, SOS2 | 0.009186 |
| hsa04926 | Relaxin signaling pathway | MMP13, GNG5, EGFR, PIK3CA, NFKBIA, JUN, CREB3L1, MMP1, SOS1, SOS2 | 0.009766 |
| hsa00230 | Purine metabolism | RRM1, APRT, POLR2J2, POLR2, GPOLE4, POLR3K, PRIM1, ATIC, POLR2C, POLR2D, GART, ENPP4, NTPCR, POLR1C, POLR2, IPOLR2J | 0.005157 |
| hsa04723 | Retrograde endocannabinoid signaling | NDUFA9, NDUFB2, NDUFB7, NDUFS1, NDUFS3, PLCB4, NDUFA3, NDUFA4, MAPK1, NDUFA8 | 0.005859 |
| hsa04060 | Cytokine-cytokine receptor interaction | CCL2, EGFR, EDA, IL2RG, CXCL10, LIF, TNFRSF11B, XCL2 | 0.007813 |
| hsa03050 | Proteasome | PSMB10, PSMC2, POMP, PSMA2, PSMA, PSMB1, PSMB4, PSMD14 | 0.007813 |
| hsa00190 | Oxidative phosphorylation | ATP5PB, ATP6V1A, COX6B2, NDUFA3, ATP6V0D1, ATP5PO, COX5B, COX6B1, COX7B, COX8A, UQCRQ, UQCRC1, NDUFA4, UQCRFS1, NDUFA8, UQCR11, NDUFA9, NDUFB2, NDUFB7, NDUFS1, NDUFS3, ATP5F1B | 0.000256 |

**Table.5.** List of pro-survival pathway genes significantly expressed in MV infected cells.

| Related GO term | Description | Gene name | Fold change | P value |
| --- | --- | --- | --- | --- |
| GO:0032956 | Actin<br>cytoskeleton<br>rearrangement | ARF6 | 2.3 | 0.002523 |
|  |  | ARHGAP28 | 11.4 | 0.00474 |
|  |  | ARHGAP40 | 17.8 | 0.000297 |
|  |  | OPHN1 | 41.7 | 0.000301 |
|  |  | TMSB4X | 0.47 | 0.003534 |
|  |  | TRIOBP | 17.4 | 0.003334 |
|  |  | CAPZB | 12.8 | 0.045622 |
|  |  | TPM3 | 2.3 | 0.041 |
|  |  | TACSTD2 | 1.7 | 0.039126 |
|  |  | MICAL3 | 23.6 | 0.005866 |
| GO:0006914 | Autophagy | GABARAPL1 | 11.3474 | 0.000154 |
|  |  | MAP1LC3B | 2.1 | 0.004651 |
|  |  | RAB7A | 9.1 | 3.45E-08 |
|  |  | FEZ1 | 29.4 | 8.21E-06 |
|  |  | LRSAM1 | 10.092 | 0.001255 |
|  |  | TICAM1 | 3.54946 | 0.00028 |
|  |  | TRIM5 | 23.5817 | 0.001425 |
|  |  | PIK3CA | 2.34366 | 0.005209 |
| GO:0045087 | Immune evasion<br>signalling process | CCL2 | 41.1639 | 0.001415 |
|  |  | CD55 | 20.2412 | 0.032958 |
|  |  | RPS27A | 0.363462 | 0.028371 |
|  |  | SAMHD1 | 0.430985 | 0.013921 |
|  |  | XCL2 | 5.22208 | 0.020581 |
|  |  | TRIM15 | 40.5199 | 0.003208 |
|  |  | MSRB1 | 0.374091 | 0.004295 |
|  |  | RBCK1 | 45.03 | 0.000856 |
| GO:0007049<br>GO:0000077<br>GO:2001022 | Regulation of cell<br>cycle and DNA<br>damage response | SYF2 | 14.1682 | 9.93E-09 |
|  |  | TRIAP1 | 0.3893 | 0.000309 |
|  |  | NABP1 | 0.257413 | 1.95E-05 |
|  |  | NUDC | 0.100511 | 0.005435 |
|  |  | NUP43 | 8.52728 | 0.000821 |
|  |  | MAD2L1 | 0.398981 | 0.000124 |
|  |  | MRE11 | 0.285434 | 0.005237 |
|  |  | CDCA5 | 0.38159 | 0.00069 |
|  |  | MNS1 | 10.2083 | 3.21E-07 |
|  |  | MRNIP | 7.66825 | 0.001941 |
|  |  | EGFR | 7.72826 | 1.09E-11 |
|  |  | SMCHD1 | 6.4 | 4.25E-06 |

**Fig.4.**
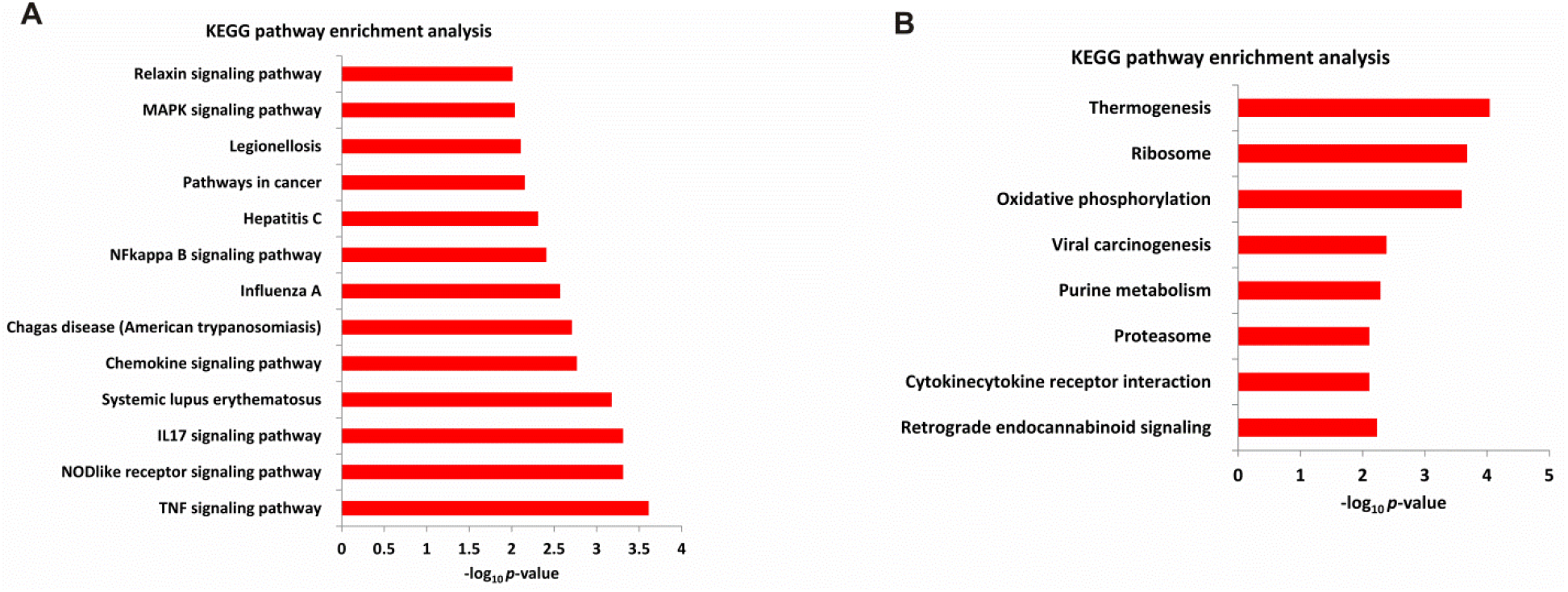
KEGG pathway enrichment analysis for genes differentially regulated in Caco-2 cells upon infection with MVs. (A) Pathways significantly regulated in 4 h post-infection. (B) Pathways significantly regulated during 8 h post-infection. Enrichment pathways were sorted based on q-value, and significance was assessed by Wilcox test.

### Endocytosis and autophagy responses

The initial attachment and binding of MVs to epithelial surface triggered a dynamic host response as observed by GO enrichment of functional groups, including response to the endocytic pathway and autophagy. Within 4 h data set, the CAV1 gene was highly up-regulated (55 fold), suggesting that MVs could be internalized via caveolin-mediated endocytosis. The transcriptome signatures also revealed modulation of cell-to-cell junction proteins, in particular, claudin 3 (CLDN3) and claudin 8 (CLDN8), which serve as the sealing components of the tight junction forming the paracellular barrier, highlighting MV invasion based on parasitosis. Transcripts involved in the autophagy process were also significantly up-regulated at 4 h infection. Importantly, GABARAPL1, FEZ1, MAPILC3B, RAB7A, ATP6V1G1 (GO:0006914) were significantly up-regulated. At 8 h infection, autophagy-related genes were not regulated considerably suggesting that the autophagy is an early stage mechanism of host response during MV infection.

### Inflammatory pathways

Gene set enrichment analyses (GSEA) revealed the up-regulation of innate immune response genes including inflammation, inflammasome signaling (encompassing Nod-like receptor and TLR7), and cytokine signaling (inflammation and cell death IL17 signaling pathway (hsa04657)). The top up-regulated genes include C-C motif chemokine ligand 2(CCL2) (FC-59.7), C-X-C motif chemokine ligand 5(CXCL5), C-X-C motif chemokine ligand 2(CXCL2), C-X-C motif chemokine ligand 3(CXCL3), C-X-C motif chemokine ligand 8(CXCL8), C-X-C motif chemokine ligand 16(CXCL16), and C-X-C motif chemokine ligand 10(CXCL10). Similarly, the transcript levels of genes encoding pro-inflammatory cytokines peaked at 4 h post infection. The up-regulated genes enriched for the KEGG-pathways include ‘cytokine-cytokine interaction pathway’ (hsa04060), NF-kappa B signaling pathway (hsa04064) ‘TNF signaling pathway’ (hsa04668) and IL-17 signaling pathways (Table 4). Expression of the pattern recognition receptors, TLRs, plays an essential role in the activation of the host immune responses. TLR7 was specifically up-regulated at 4 h of infection. At 8 h infection, up-regulated genes associated with inhibition of complement system activation (CD55, complement decay-accelerating factor), EGFR, CCL2, and XCL2 (GO:0006954). The tumor necrosis factor receptor type1 death domain (TRADD) was down-regulated, and several cytokine genes were not significantly regulated as compared to the early response.

### Response to oxidative stress and metabolic process

The induction of reactive oxygen species (ROS) by host cells represents the first line of defense against intracellular pathogens. At 4 h, the transcriptome of Caco-2 cells showed the induction of thioredoxin reductase 1 (TXNRD1), EGFR, MMP14, CFLAR, NDUFA12, ATP2A2 (GO:0006979). On the other hand, genes encoded for oxidative phosphorylation were found to be down-regulated (GO:0006119) ATP5PO, NDUFB2, COX5B, NDUFS1, UQCRQ, NDUFS3, NDUFA4, NDUFA8.

### Differential expression of genes involved in signaling pathways

Differential expression of several genes involved in signaling pathways (*P* < 0.05, Wilcox test), according to the KEGG annotation was observed. Notably, TNF signaling pathway, NOD-like receptor signaling pathway, IL-17 signaling pathway, NF-kappa B signaling pathway, MAPK signaling pathway, Relaxin signaling pathway, and PI3K-Akt signaling pathway were significantly enriched (Table 4).

### Non-coding RNAs

RNA sequencing of MV-infected host cells revealed several lincRNAs, microRNAs, snoRNAs, and snRNAs that were significantly regulated upon infection with MVs (Fig. 5). Notably, we observed 157 host snoRNAs were differentially regulated (Supplementary Table S3). These snoRNAs were categorized into 53 H/ACA box type (SNORD) and 104 H/ACA box type (SNORA) groups. Both types of snoRNAs were significantly down-regulated upon infection with MVs. In addition, long intergenic non-protein coding RNA (lincRNA) (n=10), microRNA (miRNA) (n=6), antisense RNA (n=16), small nuclear RNA 312 (snRNA) (n=13), small Cajal body-specific RNA (SCARNA1) (n=8) were significantly regulated during 4 h infection. Similarly, lincRNAs (n=4), microRNA (n=1), snoRNA (n=64), and snRNA (n=8) were significantly regulated at 8 h infection.

**Fig.5.**
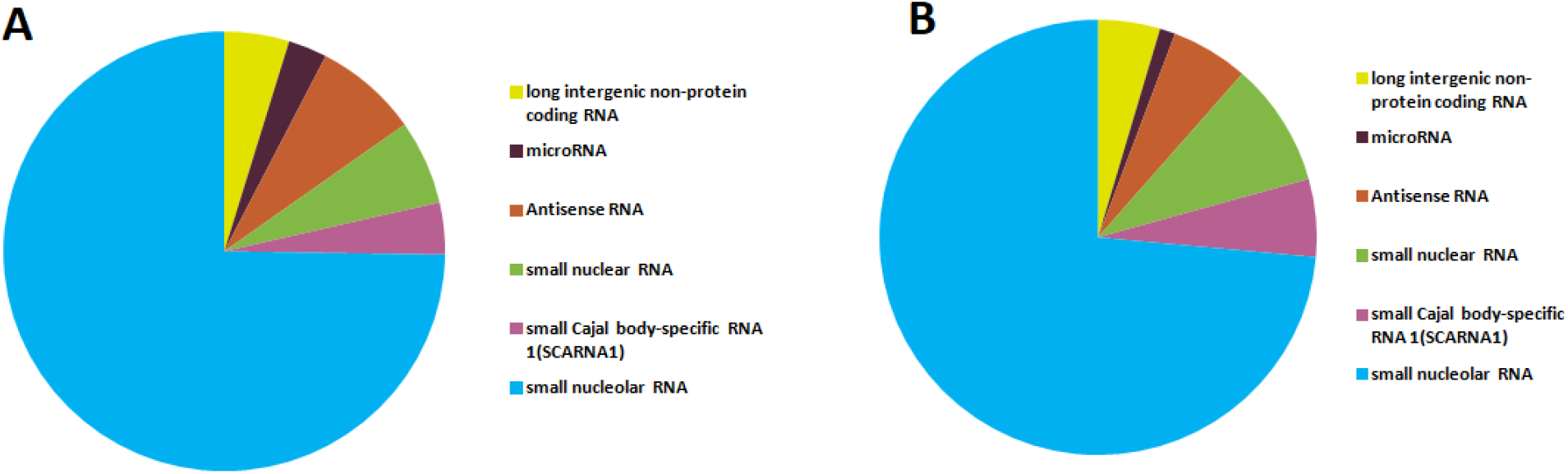
Regulation of non-coding RNAs in Caco-2 cells infected with MVs. (A) Types of differentially regulated non-coding RNAs upon infection with MVs 4 h (B) Differentially regulated non-coding RNAs in 8 h post-infection.

### Similar host responses to *L. monocytogenes* and its MVs

When compared to the uninfected cells, 2,888 genes (1,375 were up-regulated, and 1,531 were down-regulated) were differentially expressed in Caco-2 cells during 4 h post-infection, whereas 1189 (669 were up-regulated, and 520 were down-regulated) were differentially expressed in MV-infected cells (Fig. 6A). Similarly, during the 8 h infection revealed a total of 2,216 genes were significantly regulated. Of these, 727 were up-regulated, and 1,489 were down-regulated (>1.5- fold regulation, p-value<0.05, padj<0.05), whereas 989 genes were significantly regulated in MV-infected cells. Of these, 360 were up-regulated while 629 were down-regulated (>1.5- fold regulation, pvalue<0.05, padj<0.05) (Fig. 6B). Overall, 560 genes were commonly regulated in both sets during 4 h post infection wherase 523 genes were commonly regulated in both sets during 8 h post infection (Fig. 6A-B). Furthermore, we performed pathway analysis with the distinct subset of genes modulated by both treatments. The results revealed that the genes regulated by both MVs and cells showed enrichment of common GO terms. Analysis of the DE genes commonly altered by the infection of *L. monocytogenes* and MVs (Supplementary Table S4) revealed the enrichment of genes involved in the inflammatory response, cellular response to tumor necrosis factor (TNF), immune response, apoptotic cell signaling, ER stress, and others. The commonly DE genes could be categorized into several canonical pathways, such as TNF signaling, toll-like receptor (TLR) signaling pathway, nuclear Factor-kB (NF-kB), chemokine signaling, and cytokine-cytokine receptor interaction (Fig 7A). Altogether, these results suggested that both *L. monocytogenes* and its MVs could modulate the expression of a similar set of genes and pathways.

**Fig.6.**
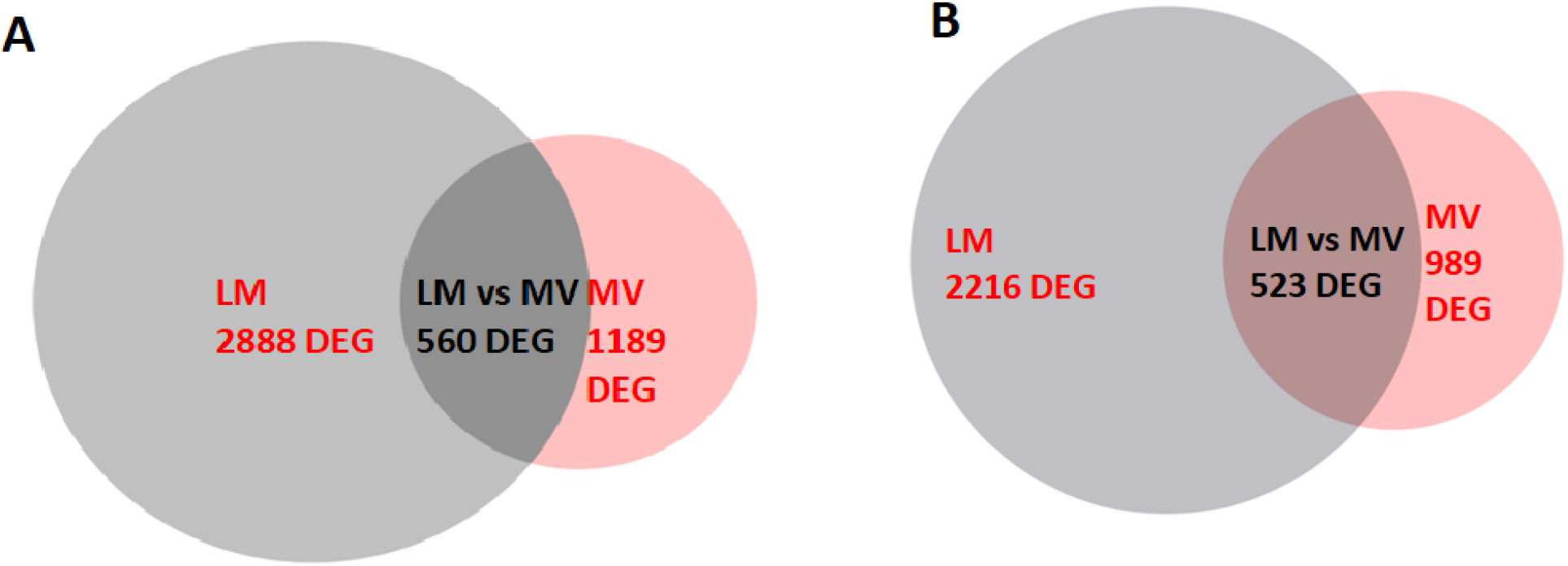
Comparison of the gene expression pattern in Caco-2 cells infected with *L. monocytogenes* and its MVs. Venn diagram depicting the number and distribution of common differentially expressed genes in 4 h (A) and 8 h post-infection (B).

**Fig.7.**
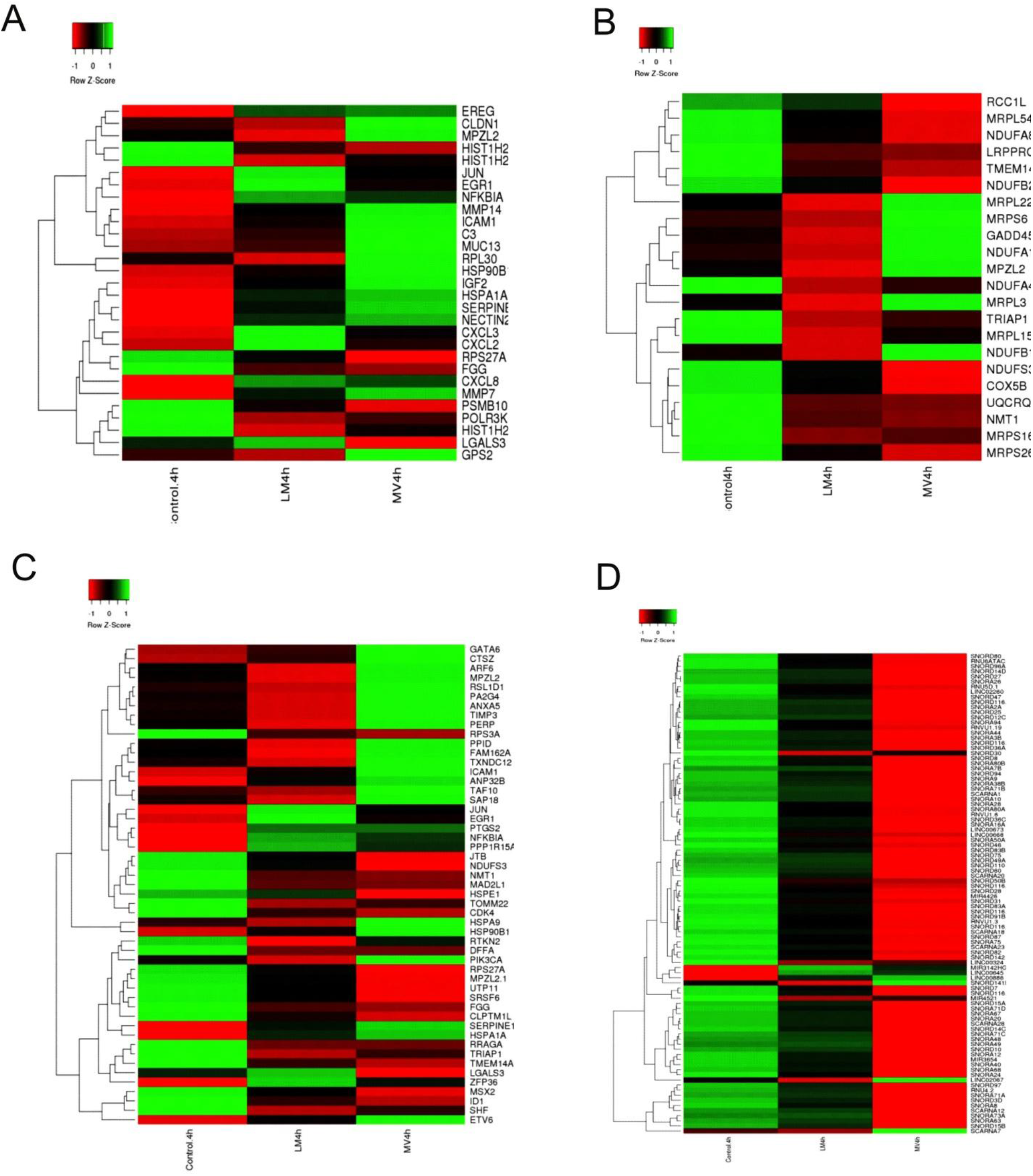
Heatmap showing differential gene expression in Caco-2 cells at 4 h of infection with *L. monocytogenes* and its MVs. Gene ontology (GO) group of the innate immune response (A), mitochondrial (B), apoptosis (C) and non-coding RNAs (D).

### Dissimilar host responses to *L. monocytogenes* and its MVs

At the early stage of infection, specific genes involved in mitochondrial-mediated transport, apoptosis, immune response, and miRNAs showed differentially regulated by *L. monocytogenes* and its MVs. For instance, mitochondrial-mediated transport genes MRPL22, MRPS6, MPZL2, NDUFS3, and COX5B were up-regulated by MVs and down-regulated by *L. monocytogenes* (Fig. 7B). Similarly, innate immune response and apoptotic related genes CXCL2, CXCL8, ICAM1, MUC13, C13, NECTIN2, GATA6, ARF3, TIMP3, NDUF3, MSX2, NFKBIA, and RPS27A were differentially regulated (Fig 7A & C). Also, several lincRNAs, microRNAs, snoRNAs, and snRNAs were significantly down-regulated by the infection with MVs when compared to *L. monocytogenes* (Fig. 7D).

At 8 h post-infection, innate immune response genes such as NFKBIA, HIST1H2BG, HIST1H2BE, ID2 were differentially regulated (Fig 8A). Similarly, the cytoskeleton and microtubule-associated genes were differentially regulated (Fig. 8B). Few cell cycle-related genes such as MCM6, MYC, and CCNB1 were up-regulated by MVs and down-regulated by *L. monocytogenes* (Fig. 8C). Likewise, mitochondrial-mediated transport genes and several lincRNAs, microRNAs, snoRNAs, and snRNAs were differentially regulated (Fig. 8 D-E).

**Fig.8.**
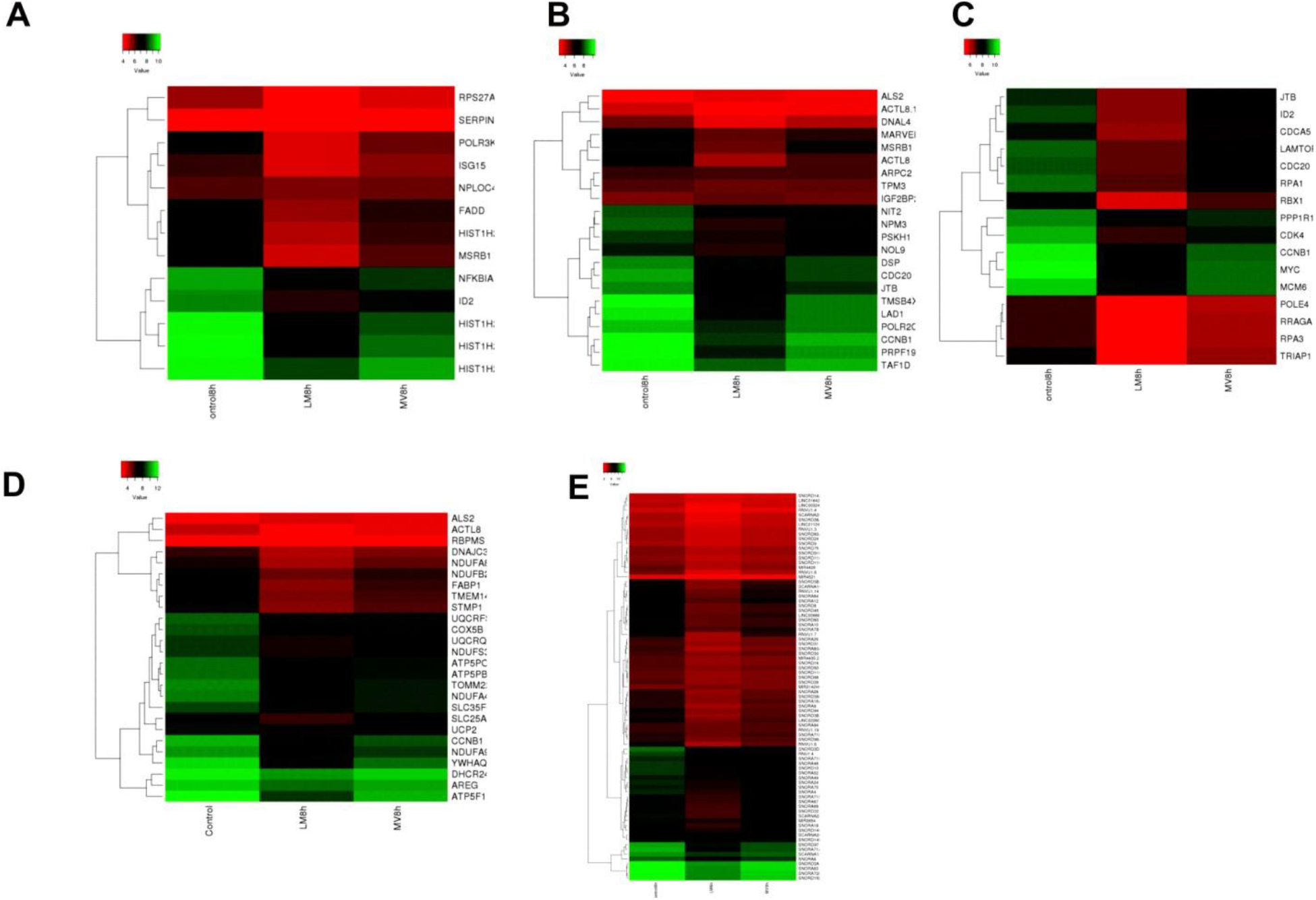
Heatmap showing differential gene expression in Caco-2 cells at 8 h of infection with *L. monocytogenes* and its MVs. Gene ontology (GO) group of the innate immune response (A), actin-cytoskeleton (B), cell cycle (C), mitochondrial (D) and non-coding RNAs (E).

Overall, 597 genes were differentially expressed when infected with MVs but not with the cells. These genes showed both positive and negative regulation of inflammatory response, endocytosis, positive regulation of apoptosis. MV-regulated genes could be categorized into several pathways, such as PI3k-Akt signaling pathway, mitogen-activated protein kinase (MAPK), NOD-like receptor signaling pathway, cAMP signaling pathway, TNF, and NF-kB signaling.

### MVs modulated pro-survival pathways of the host cell

#### (i) actin cytoskeleton rearrangements in host cells

Several genes associated with actin cytoskeletal rearrangements were up- and down-regulated. Enrichment analysis suggested that MVs induced the disruption and remodeling of the cytoskeleton. In particular, actin filament organization (GO:0007015), regulation of actin filament (GO:0030832), regulation of actin polymerization or depolymerization (GO:0008064) were within the enrichment category by GO analysis. Subsets of genes associated with actin cytoskeleton rearrangements were consistently up-regulated by MVs (Fig. 9). Importantly, the process related to actin cytoskeleton rearrangement and vacuolization were the most enriched within the up-regulated genes. In particular, ARHGAP28, ARHGAP40, ARF, TRIOBP, and OPHN1 were up-regulated at 4 h, and CAPZB, TPM3, TRIOBP, TACSTD2, TMSB4X, and MICAL3 were up-regulated at 8 h.

**Fig.9.**
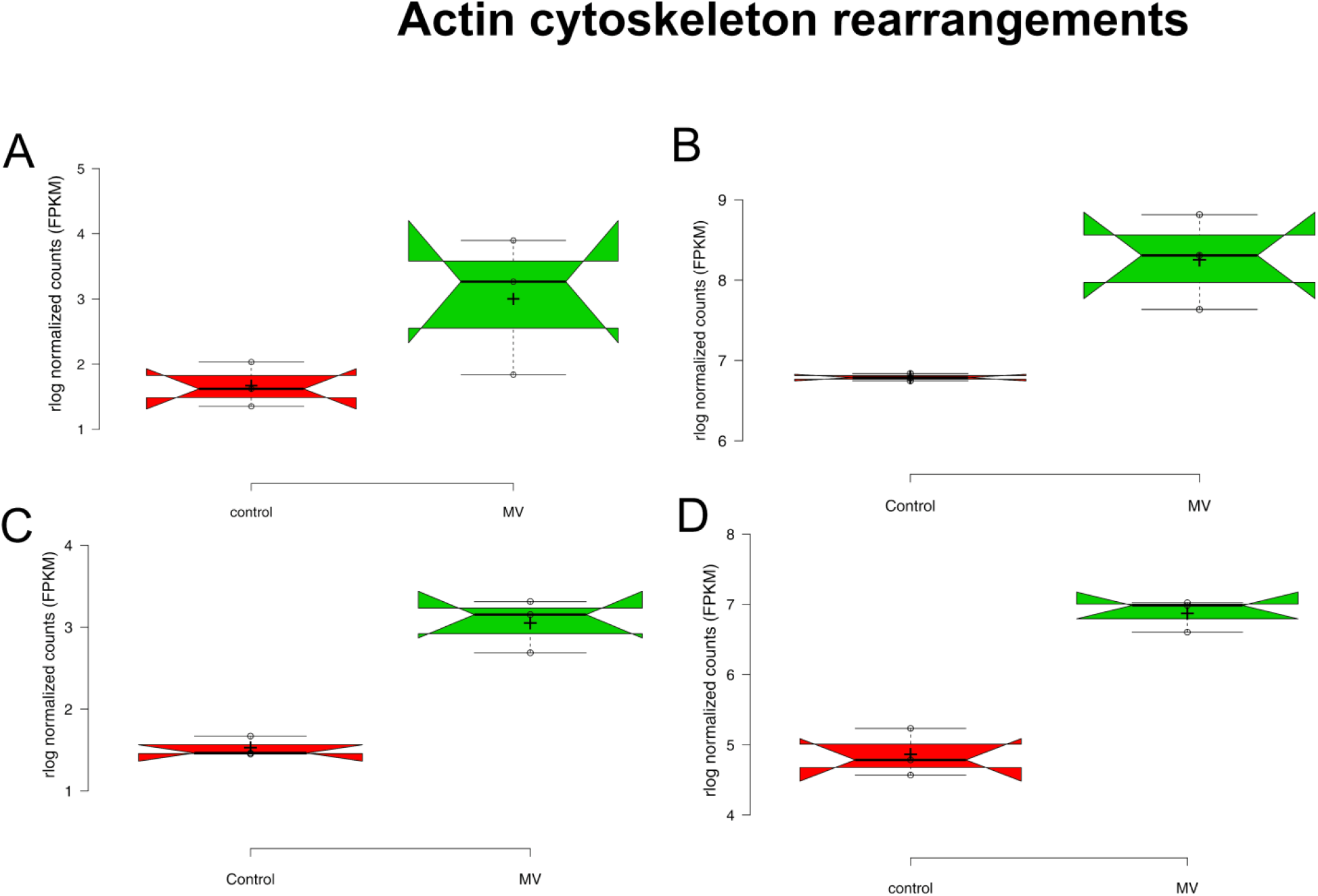
Differentially expressed genes belong to actin-cytoskeleton rearrangement (GO:0032956) in response to MV infection. (A) ARF (B) ARHGAP28 (C) OPHN1(D) WASF2. Red notch indicates uninfected control and green indicate test.

#### (ii) Autophagy and xenophagy response

GO term analysis revealed that autophagy-related genes were significantly up-regulated during the early response by MVs and down-regulated by *L. monocytogenes* cells. Autophagy and xenophagy (macroautophagy) related genes, including GABARAPL1, FEZ1, MAP1LC3B, and RAB7, were significantly up-regulated by MVs in early response (Fig. 10). At 8 h infection, only GABARAPL1 gene was found to be differentially expressed, whereas other autophagy genes were not significantly regulated, suggesting a dramatic remodeling of autophagosome formation following MV infection. Thus, MVs could inhibit or avoid autophagy of the host cells to favor the intracellular survival of *L. monocytogenes*.

**Fig. 10.**
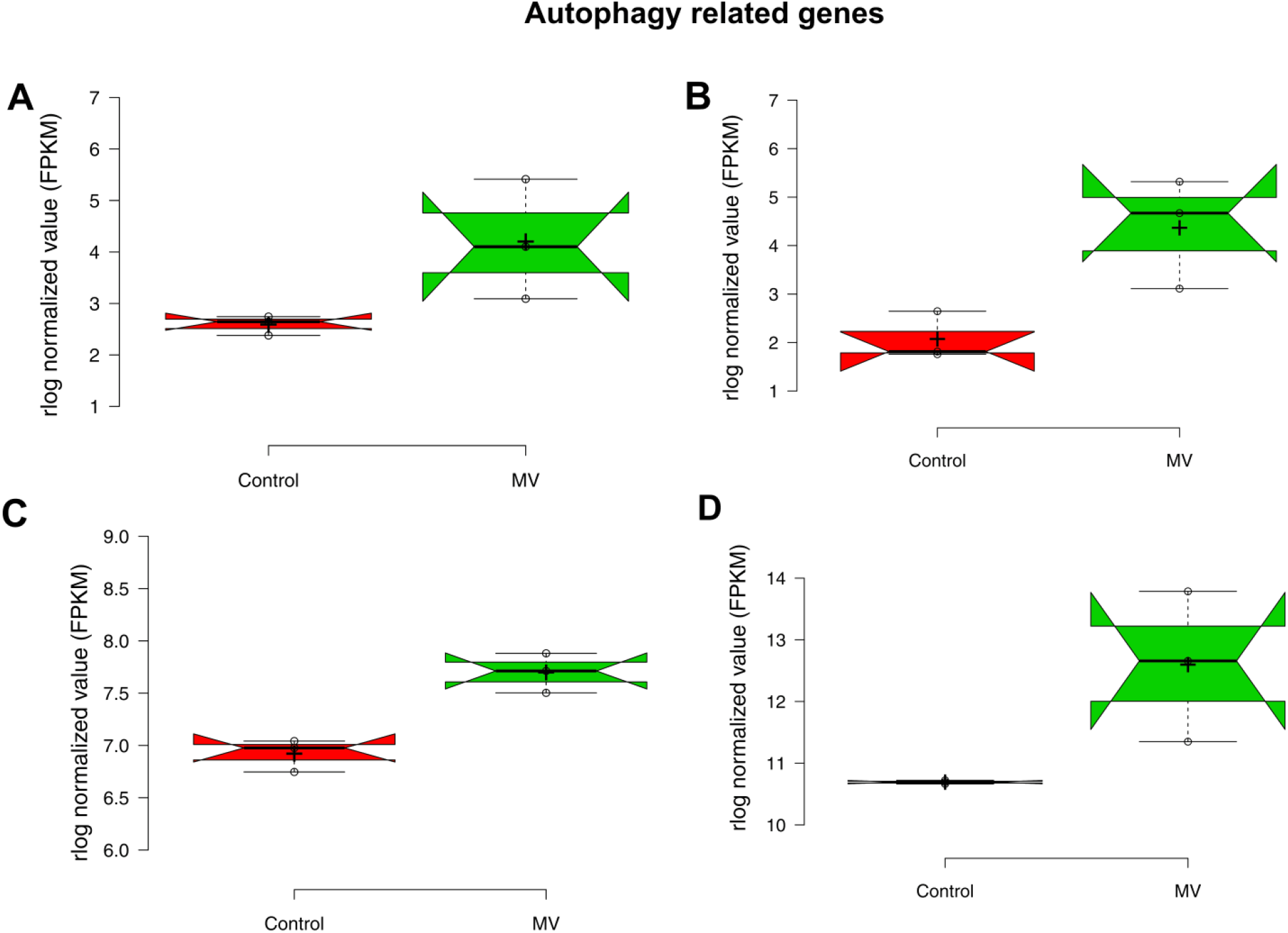
Differentially expressed genes belong to autophagy (GO:0006914) in response to MV infection. (A) GABARAPL1 (B) FEZ1 (C) MAP1LC3B (D) RAB7A. Red notch box indicates uninfected control and green indicate test.

#### (iii) Immune evasion signals

At 4 h infection, many cytokine-related genes were significantly up-regulated (Fig. 11). However, only a few cytokines CCL2, XCL2, CXCL10 were up-regulated considerably at 8 h. Notably, overexpression of TRIM15 and RBCK1 was observed at 8 h. The up-regulation of RBCK1 can negatively regulate TAB2/3 and TNF induced NF-KB activation. Also, overexpression of RBCK1 can inhibit the inflammatory signaling cascade.

**Fig. 11.**
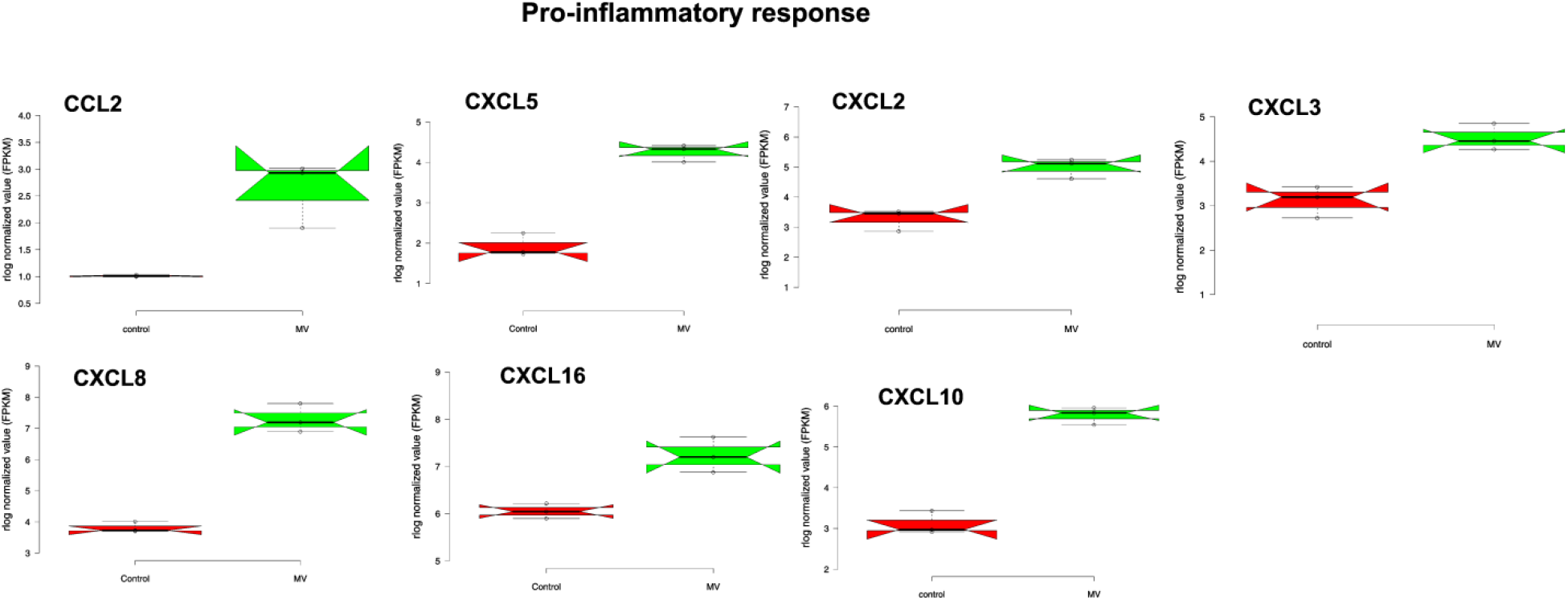
Differentially expressed pro-inflammatory response genes in response to MV infection. The top up-regulated genes were those encoding the C-C motif chemokine ligand 2(CCL2), C-X-C motif chemokine ligand 5(CXCL5), C-X-C motif chemokine ligand 2(CXCL2), C-X-C motif chemokine ligand 3(CXCL3), C-X-C motif chemokine ligand 8(CXCL8), C-X-C motif chemokine ligand 16(CXCL16), C-X-C motif chemokine ligand 10(CXCL10). The red notch box indicates uninfected control and green indicate test.

#### (iv) Regulation of cell cycle and DNA damage response

Oxidative stress induces DNA damage, which if not fixed, leads to cell cycle arrest and apoptosis. MVs differentially regulated cell cycle associated genes SYF2, TRIAP1, NABP1, NUDC, NUP43, and MAD2L1. The checkpoint gene MRE11 was significantly down-regulated (Fig. 12) during the early infection, suggesting a positive regulation of the cell cycle. Also, the DNA damage response genes such as MRNIP, MNS1, EGFR, and SMCHD1 were significantly up-regulated during MV infection, suggesting a dramatic response to DNA damage followed by MV infection.

**Fig.12.**
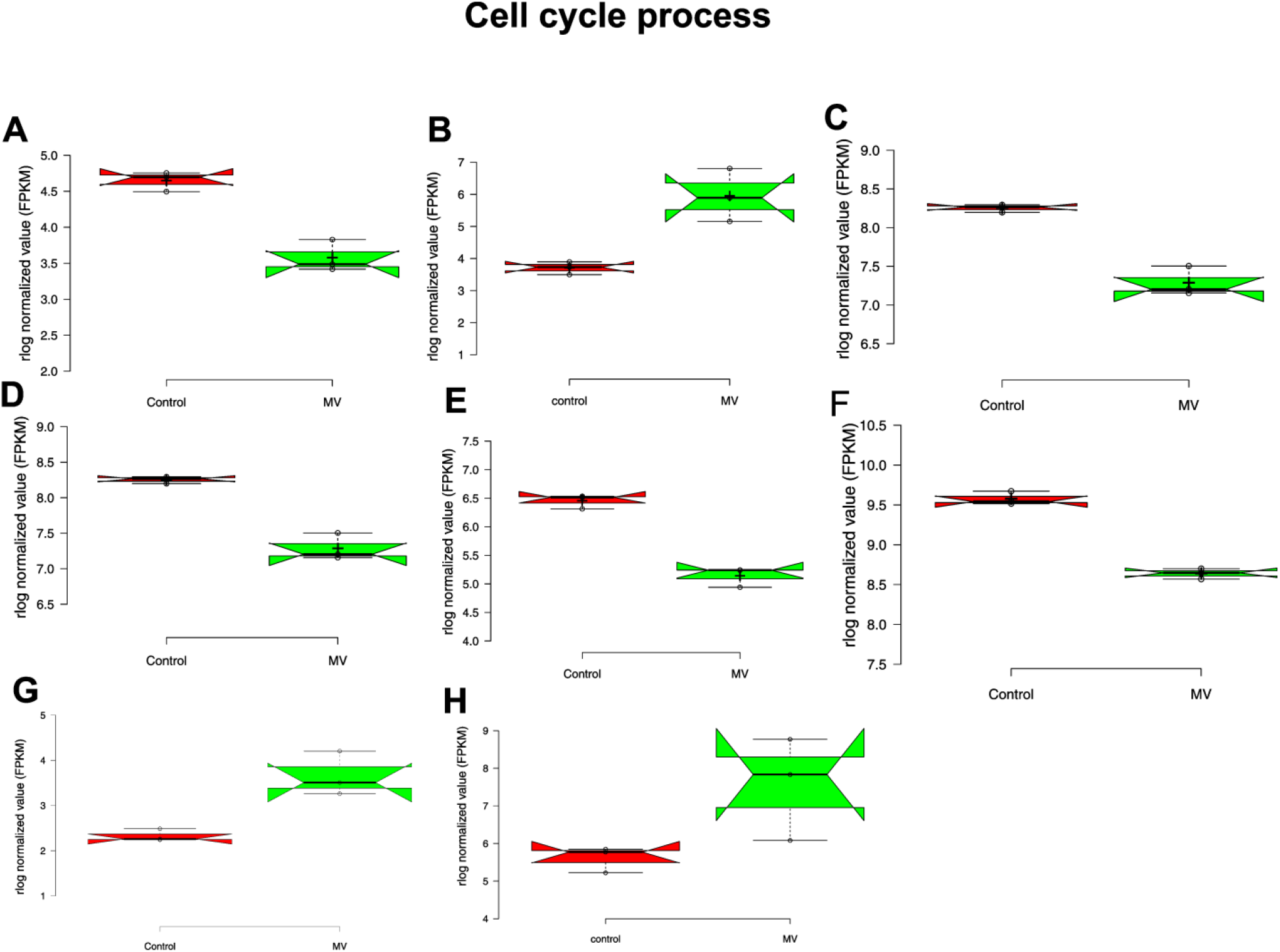
Differentially expressed genes belong to the regulation of cell cycle and DNA damage response (GO:0007049), (GO:0000077), GO:2001022: (A) MRE1, (B) SYSF2 (C) CDCA5 (D) TRIAP1 (E) NABP1 (F) MAD2L1 (G) MRNIP, (H) MNS. Red notch indicates uninfected control and green indicate test.

## Discussions

Secretion of outer-membrane vesicles is a fundamental process for both Gram-positive and Gram-negative bacteria. These outer membrane vesicles are mainly involved in the transfer of DNA, proteins, cell to cell signaling, biofilm formation, stress response, delivery of toxin to the host cell and host cell interactions (8). MVs of Gram-negative bacteria and their roles in virulence are well characterized (6–8). Recent data suggest that vesicles of Gram-positive bacteria also shown to involve in virulence. Previous studies from several Gram-positive pathogenic bacteria including, *S. aureus* (34,35), *Streptococcus suis* (36), *Bacillus anthracis* (11), Group B Streptococcus (37), *Mycobacterium tubercul*osis (38), *Clostridium perfringens* (13) secretes biologically active MVs and known to involve in pathogenesis. These studies suggested that MVs derived from Gram-positive play an essential role in bacterial pathogenesis as similar to MVs from Gram-negative bacteria.

However, very little information is available on the functional characterization of MVs of *L. monocytogenes* (14,39). Here, we studied the host cellular responses to the MVs of *L. monocytogenes* and compared the responses to the bacterial cells. Proteome analysis of MVs of *L. monocytogenes* revealed that 456 proteins are derived from the cytoplasm, membrane and cell wall, and localization of few proteins could not be predicted. The majority of them were predicted to be cytoplasmic proteins. Recent studies are suggesting that MVs are naturally incorporating a large number of cytoplasmic proteins into MVs. The mechanism by which these cytoplasmic proteins are differentially packaged in the MVs is unknown. Many of the identified cytoplasmic proteins could be moonlighting proteins at the cell surface of *L. monocytogenes*. This group includes many conserved proteins involved in central metabolic pathways, cellular responses to stress, and/or virulence.

We have also identified several lipoproteins in MVs. In Gram-positive bacteria, lipoproteins are involved in nutrient transport, Toll-like receptor 2 activations, and pathogenicity (40). A recent study has demonstrated that lipoproteins are essential for the virulence of *L. monocytogenes* (41). The proteomic profiling of MVs in this study provides a better understanding of the selective sorting of virulence factors and the general physiological attributes of MVs and the MV-mediated virulence mechanisms in *L. monocytogenes*. Also, we identified several chaperone proteins associated with MVs. These proteins were reported to be involved in the virulence and stress response (42). We have identified major *Listeria* transporter proteins in MVs such as OpuCA, OpuCC and it has been reported that loss of OpuC reduces the intestinal colonization of *L. monocytogenes* during infection (43).

In this study, we selected the human epithelial cell line Caco-2 cells because it has been successfully used as an infection model to investigate the pathogenesis of *L. monocytogenes* and other bacteria. Recently, our group characterized the interaction of MVs with Caco-2 cells and demonstrated that MVs were internalized via actin-mediated endocytosis (19). However, despite an increase in our understanding of the pathogenic mechanism and immune response to *L. monocytogenes* and its MVs, the host cell response to the MVs remains unclear. Thus, the response of Caco-2 cells to *L. monocytogenes* derived MVs were determined using RNAseq. The GO enrichment analysis in biological processes revealed the unique features of MV infection. For early up-regulated groups, the most enriched cluster was those that participate in endocytosis, defense response, and pro-inflammatory response. In contrast, the most enriched early down-regulated group were involved in the regulation of metabolic pathways. A previous study has reported the transcriptome of Caco-2 cells infected with *L. monocytogenes* by microarray. They found that the innate immune regulatory genes are actively regulated upon infection with *L. monocytogenes* (44).

The major events occurring during the interplay between MVs and the host cells were identified in our transcriptome analysis. We found that MVs substantially regulate the actin cytoskeleton network, leading to rearrangement or vacuolization, as highlighted by the GO enrichment category. Loss of cytoskeletal integrity and increased epithelial permeability are usually hallmarks of inflammation. Cytoskeletal rearrangement promotes numerous events that are beneficial to the pathogen, including internalization of bacteria, structural support for bacteria-containing vacuoles, altered vesicular trafficking, actin-dependent bacterial movement, and pathogen dissemination (45). Also, MVs could trigger vacuolization for *L. monocytogenes* to survive or persist inside the vacuoles during infection. This observation in agreement with the previous study demonstrated that *L*. *monocytogenes* switches from this active motile lifestyle to a stage of persistence in vacuoles (46). Other studies have also demonstrated that OMVs enhance the intracellular vacuole formation during infection with *Legionella pneumophila* (7). Further, autophagy-related genes were significantly up-regulated in response to the early infection of MVs, and not expressed during the late phase of infection. Thus, MVs could induce autophagy in the early stage of infection, which is in agreement with a previous study demonstrated that the MVs from *L. monocytogenes* inhibit autophagy (39).

Recognition of infectious agents by host cells results in the modulation of transcriptional programs to tackle the infection (47). However, the activation of innate immunity by pattern recognition receptors (PRRs) in response to infection with *L. monocytogenes* is still poorly understood. Also, pattern recognition receptor that mediates recognition of MVs is still unknown. In this study, TLR7 was up-regulated following the infection with MVs. Generally, the TLR family recognize various categories of pathogen-associated molecules. Peptidoglycan from *L. monocytogenes* is recognized by TLR2 (48) and significant virulence factor listeriolysin O is a ligand for TLR4 (49). Recently, TLR10 was also regulated in macrophage as well as epithelial cells upon infection with *L. monocytogenes* (50). Here, we found that TLR7 recognizes MVs of *L. monocytogenes*. However, possible ligand from the MVs is unknown. The murine TLR7 protein (human orthologue, TLR7/8) is shown to respond to the viral single-stranded RNAs (ssRNAs), as well as to Streptococcal bacterial RNAs in dendritic cells (51,52). Possibly, RNAs packaged in the MVs could activate host signal cascades via TLR7.

Numerous genes involved in the innate and adaptive immune responses were regulated in response to MV infection. The component of the innate immune response such as CCL2, CXCL8, CXCL5, CXCL16 was among the most up-regulated genes, highlighting the recruitment of T cells and macrophages, a phenomenon associated with inflammation. However, at 8 h post-infection, these genes were not significantly regulated, possibly to avoid excessive inflammation. Modulation of immune signals in the host has been demonstrated as an essential strategy developed by successful pathogens to subvert host responses, switching the immune responses into a hypo-responsive state (53). Regulation of expression of genes associated with different signaling cascades such as TLR, TNFR, NF-kB and Akt signaling pathway in the MV-infected cells suggests that the MVs may modulate innate immune system components at various levels to evade and subvert host responses. Previous studies have demonstrated that the innate immune response is activated in response to *L. monocytogenes* infection in Caco-2 cells (44).

A strategy developed by the intracellular pathogens to establish a niche of infection is the ability to control host cell damage and cell cycle to their advantage. MVs promoted increased expression of cell cycle regulatory genes such as cell cyclins and CDK, which may enhance host cell survival of *L. monocytogenes*. Notably, down-regulation of DNA damage response gene MRE11 was observed. These results corroborate with previous reports demonstrated that listeriolysin O could degrade mre11 and promote bacterial replication (54). Thus, the listeriolysin O enriched in MVs may interact with host cell to promote dysregulation of DNA damage response. Similarly, the up-regulation of cell cycle-related genes in response to MV could also be a strategy to promote host cell survival, in turn, to retain a replicative niche for *L. monocytogenes*.

Another important observation in this study is the regulation of lincRNAs, microRNAs, snoRNAs, and snRNAs upon infection with MVs. Survival of intracellular pathogens in host cells depends on the modulation of several cellular functions by regulating the host gene expression (47). Manipulation of the host transcriptome serves as a selective advantage for the intracellular life cycle of the pathogen. Our findings suggest that MVs not only appears to interfere with cellular functions but promoting changes in several transcriptional and posttranscriptional regulatory elements that may extensively impact host cell functions during the later stage of infection. One such example is the differential expression of several non-coding RNAs, which have regulatory roles in physiological and pathological responses (55). Down-regulation of several ncRNAs upon infection with MVs also raise interesting questions on the role of this class of RNAs in *L. monocytogenes* infection. Several reports on ncRNAs suggested that the regulatory RNAs used by intracellular pathogens may aid to survive and evade immune responses (56,57).

Overall, this study represents the first transcriptome profiles of the host cells infected with MVs of *L. monocytogenes*. The RNA-Seq data revealed the dynamic changes of cellular pathways during the infection and provided cues for understating the MV-induced host response. In particular, at late stages of infection, when bacteria are mostly paracellular or internalized, the shift in lifestyle leads to the down-regulation of the central metabolism as found in this study. The findings have opened the way for more detailed studies on the roles of MVs in the host-pathogen interaction during *L. monocytogenes* infection.

## Supporting information

Table S1. List of identified cellular proteins and peptides.xlsx

Table S2. List of identified MV proteins and peptides.xlsx

Table S3. List of differentially regulated non-coding RNAs

Table S4. List of differentially expressed genes in both L. monocytogenes and MV infection

## ACKNOWLEDGMENTS

The authors gratefully acknowledge the University Grants Commission, New Delhi, India, for providing financial support (UGC-MRP 41-1143/2012(SR)). RK thanks UGC-BSR for awarding UGC-Meritorious Fellowship. RK thanks Mr. T. Elanthendral for his support in RNA-Seq data analysis. We acknowledge the Proteomics Facility at CCMB, Hyderabad. We also acknowledge the UGC-CAS, NRCBS, DBT-IPLS, and DST-PURSE Programs of the School of Biological Sciences, Madurai Kamaraj University.

## Supplementary files

**Table S1. List of identified cellular proteins and peptides.xlsx**

**Table S2. List of identified MV proteins and peptides.xlsx**

**Table S3. List of differentially regulated non-coding RNAs**

**Table S4. List of differentially expressed genes in both *L. monocytogenes* and MV infection**

